# An improved encoding of genetic variation in a Burrows-Wheeler transform

**DOI:** 10.1101/658716

**Authors:** Thomas Büchler, Enno Ohlebusch

## Abstract

**Motivation:** In resequencing experiments, a high-throughput sequencer produces DNA-fragments (called reads) and each read is then mapped to the locus in a reference genome at which it fits best. Currently dominant read mappers (Li and Durbin, 2009; Langmead and Salzberg, 2012) are based on the Burrows-Wheeler transform (BWT). A read can be mapped correctly if it is similar enough to a substring of the reference genome. However, since the reference genome does not represent all known variations, read mapping tends to be biased towards the reference and mapping errors may thus occur. To cope with this problem, Huang *et al*. (2013) encoded SNPs in a BWT by the IUPAC nucleotide code (Cornish-Bowden, 1985). In a different approach, Maciuca *et al*. (2016) provided a ‘natural encoding’ of SNPs and other genetic variations in a BWT. However, their encoding resulted in a significantly increased alphabet size (the modified alphabet can have millions of new symbols, which usually implies a loss of efficiency). Moreover, the two approaches do not handle all known kinds of variation.

**Results:** In this article, we propose a method that is able to encode many kinds of genetic variation (SNPs, MNPs, indels, duplications, transpositions, inversions, and copy-number variation) in a BWT. It takes the best of both worlds: SNPs are encoded by the IUPAC nucleotide code as in (Huang *et al*., 2013) and the encoding of the other kinds of genetic variation relies on the idea introduced in (Maciuca *et al*., 2016). In contrast to Maciuca *et al*. (2016), however, we use only one additional symbol. This symbol marks variant sites in a chromosome *and* delimits multiple variants, which are added at the end of the ‘marked chromosome’. We show how the backward search algorithm, which is used in BWT-based read mappers, can be modified in such a way that it can cope with the genetic variation encoded in the BWT. We implemented our method and compared it to BWBBLE (Huang *et al*., 2013) and gramtools (Maciuca *et al*., 2016).

**Availability:** https://www.uni-ulm.de/in/theo/research/seqana/

## 1 Introduction

Low-cost genome sequencing gives unprecedented information about the genetic structure of populations. The genetic content of a population or species is often represented by a reference genome (in form of a DNA sequence for each chromosome) and a catalog of variations. A prime example is the human species. In the first draft of the human reference genome (Lander *et al*., 2001), variable regions were poorly represented. In 2008, The 1000 Genomes Project Consortium (2015) started a project to produce a catalog of all variations in the human population. Its original goal was to sequence the genomes of at least 1000 humans from all over the world. From the 2504 individuals characterized by the 1000 Genomes Project, it is estimated that the average diploid human genome has around 4.1–5 million point variants such as single nucleotide polymorphisms (SNPs), multi-nucleotide polymorphisms (MNPs), and short insertions or deletions (indels). Moreover, it carries between 2100 and 2500 larger structural variants (SVs) such as large deletions, duplications, copy-number variation, inversions, and translocations (The 1000 Genomes Project Consortium, 2015).

Since *de novo* assembly of e.g. mammalian genomes is still a serious problem (both from a technological and a budgetary point of view), the reference-based approach is usually used to detect genetic variations. In this approach, a high quality genome assembly of a single selected individual (or a mixture of several individuals) is produced, and this assembly is used as a reference for genomes in the population. Such a linear reference provides a coordinate system: the location of a genetic element is defined by its coordinate (starting position) in the reference genome. In resequencing experiments, a high-throughput sequencer produces DNA-fragments (called reads) of a certain length (which depends on the technology used) and each read is then mapped to the locus in the reference genome at which it fits best. A read can be mapped correctly if it is similar enough to a substring of the reference genome. However, since the reference genome does not represent all known variations, read mapping tends to be biased towards the reference and mapping errors may thus occur. Consequently, read mappers should address this problem by taking known genetic variations (e.g. given in a VCF-file) into account. In the following, we will show how this can be achieved for BWT-based read mappers.

### 1.1 Background

In this section, we briefly introduce the data structures on which our new algorithms are based. For details, we refer to the textbook (Ohlebusch, 2013), and the references therein.

Let Σ be an ordered alphabet of size *σ* whose smallest element is the sentinel character $. In the following, *S* is a string of length *n* on Σ having the sentinel at the end (and nowhere else).

For 1 ≤ *i* ≤ *n, S*[*i*] denotes the *character at position i* in *S*. For *i* ≤ *j, S*[*i..j*] denotes the *sub-string* of *S* starting with the character at position *i* and ending with the character at position *j*. Furthermore, *S_i_* denotes the *i*-th suffix *S*[*i..n*] of *S*. The *suffix array* SA of the string *S* is an array of integers in the range 1 to *n* specifying the lexicographic ordering of the *n* suffixes of *S*, that is, it satisfies *S*_sA[1]_ < *S*_sA[2]_ < ⋯ < *S*_sA[1]_; see Table 1 for an example. A suffix array can be constructed in linear time; see e.g. (Puglisi *et al*., 2007). For every substring *ω* of *S*, the *ω*-interval is the suffix array interval [*i..j*] so that *ω* is a prefix of *S*_sA[*k*]_ if and only if *i* ≤ *k* ≤ *j*.

**Table 1:**
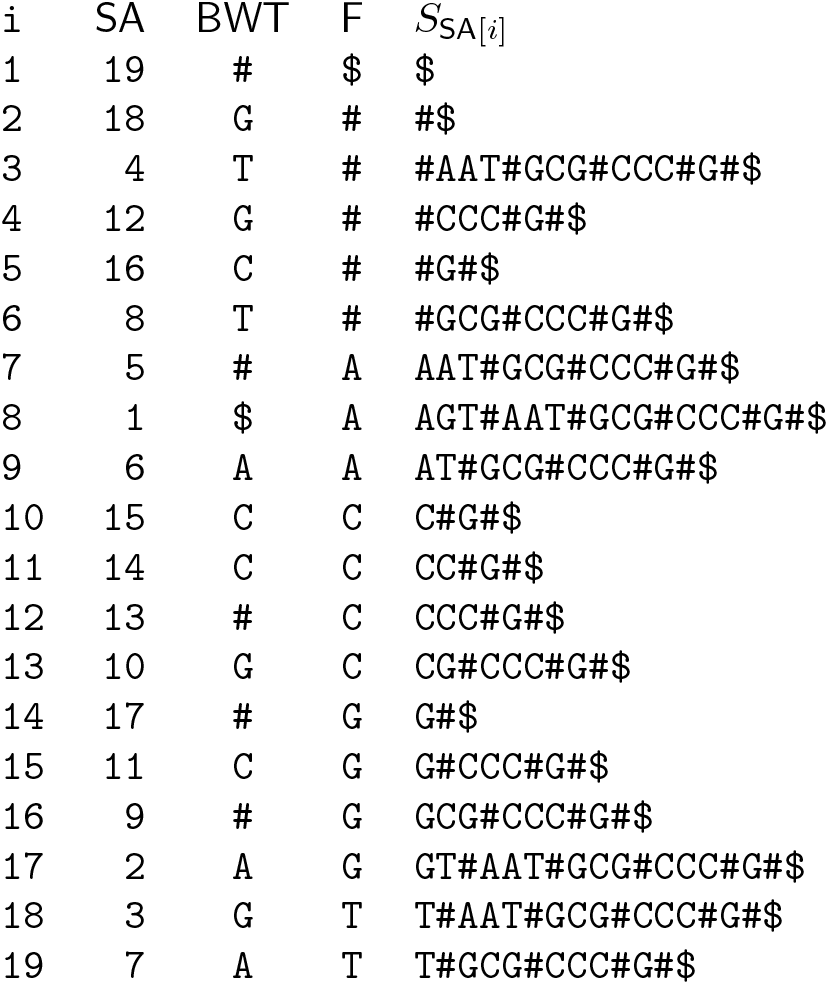
Suffix array SA and BWT of *S* = **AGT#AAT#GCG#CCC#G#$**. The columns F and *S*_sA[*i*]_ are not part of the index data structure.

The Burrows-Wheeler transform (Burrows and Wheeler, 1994) converts *S* into the string BWT[1..*n*] defined by BWT[*i*] = S[SA[*i*] − 1] for all *i* with SA[*i*] ≠ 1 and BWT[*i*] = $ otherwise.

Using the C-array (for each *c* ∈ Σ, C[*c*] is the overall number of occurrences of characters in BWT that are strictly smaller than *c*) and a data structure that supports rank-queries in the BWT (*rank*_BWT_(*i, c*) asks for the number of occurrences of character *c* in the BWT up to but excluding position *i*), it is possible to search backwards for a pattern in *S* (Ferragina and Manzini, 2000): Given an *ω*-interval [*i..j*] and *c* ∈ Σ, the procedure call *backwardSearch*(*c*, [*i..j*]) returns the *cω*-interval [*1..r*], where *l* = C[*c*]+ *rank*_BWT_(*i, c*) + 1 and *r* = C[*c*] + *rank*_BWT_(*j* + 1, *c*)]. If *l* > *r*, then *cω* is not a substring of *S*. In the computer science literature, a data structure that supports backward search in *S* is often called FM-index of *S*. In our implementation, we use a wavelet tree (Grossi *et al*., 2003), which supports rank-queries in *O*(log *σ*) time, as FM-index.

## 2 Methods

In this article, the term ‘pan-genome’ refers to the known genomic content of a certain population or species, while a ‘pan-genome index’ is a data structure that represents a pan-genome and supports efficient search within the pan-genome. The genomes of individuals of the same species are usually very similar. Thus, the DNA sequence of a chromosome can be viewed as a sequence of conserved regions (substrings common to all individuals of a population) interspersed with variant sites at which different alternatives are possible; see Figure 1 for a toy example. In this section, it will be shown how to use this model to encode genetic variants in a BWT, but first we discuss related work.

**Figure 1:**
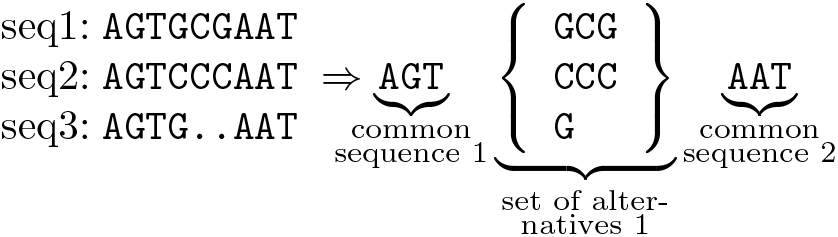
Three DNA-sequences that share **AGT** at the beginning and **AAT** at the end. The sequences differ in the middle.

### 2.1 Related work

#### 2.1.1 Encoding in *BWBBLE*

Huang *et al*. (2013) used a reference genome in conjunction with genomic variant information. They treat SNPs differently from other variants. SNPs are the most common type of sequence variation, in which the set of alternatives at a variant site consists solely of single nucleotides. As already mentioned, Huang *et al*. (2013) encoded SNPs by the 16-letter IUPAC nucleotide code (Cornish-Bowden, 1985). A IUPAC character can represent all the nucleotides that have been observed at the same position in the sequenced genomes. For example, the letter W encodes A and T. If an A should be matched in a backward search, the algorithm must also consider the letter W and all other IUPAC characters that encode A.

An indel variant at a specific locus is appended at the end of the reference genome, where a special symbol # delimits the variants. To facilitate search, each appended indel variant is padded at both ends with the bases surrounding the locus in the reference genome. The length of the padding depends on the expected read length. It must be long enough because the search algorithm must be able to map a read (containing an indel) to one of the appended sequences. Therefore, the backward search is limited to small enough patterns. Moreover, combinations of nearby variations are not considered. It is also described in (Huang *et al*., 2013) how inversions, translocations, and duplications can be handled similarly, but it seems that this is not implemented in *BWBBLE*. A major drawback of this approach is that the appended sequences significantly increase the size of the string for which an index must be built.

#### 2.1.2 Encoding in *gramtools*

Maciuca *et al*. (2016) developed a method that places the set of alternatives directly at the variant site at which they appear. In their encoding, each variant site (including SNPs) is assigned two unique numeric identifiers, one even and one odd, which they call variation markers. The odd identifiers mark variant site boundaries and the alternatives appear between these site boundaries. The even identifiers serve as separators between the alternatives. If an odd identifier is encountered in a backward search, the suffix array intervals corresponding to the alternatives must be considered; see (Maciuca *et al*., 2016) for details. It is a significant disadvantage of this approach that the alphabet size increases proportionally to the number of variant sites.

### 2.2 Encoding in *jisearch*

The method to encode SNPs by the IUPAC code is a very efficient way to deal with the vast majority of genetic variants. Thus our software tool *jisearch* (for *jump index search*) encodes SNPs in the same way. In contrast to previous approaches, however, we allow a set of alternatives to occur at different variant sites. This enables the possibility of encoding structural variants such as duplications and transpositions. Furthermore, we merely add one new symbol # to the alphabet. If # is encountered in a backward search, this results in a ‘jump’. Information about the jumps is maintained in a data structure called *jump index array* and the overall index is called *jump index*. The string containing the sequence data is called *jump index string*. It consists of the marked chromosome (which is obtained by marking every variant site with a #), followed by a list of all alternatives that occur at the variant sites. The alternatives are separated by # as well; see Figure 2 for an example. We differentiate between the two functions of a marker. If it marks a variant site, we will call it a site marker. If it separates alternatives, we will call it separator. Although all markers are encoded by the same symbol #, we can distinguish them by their rank (number of occurrences) in the *jump index string*. In Figure 2, the first occurrence of # marks a variant site while all other markers are separators. Both site markers and separators will be used as sources and targets of jumps.

**Figure 2:**
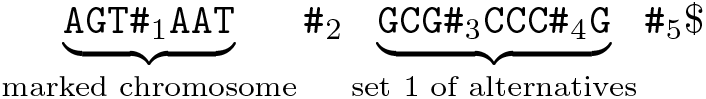
The *jump index string* of the example from Figure 1. The indices of the markers show their rank (they are numbered consecutively).

During the construction of our index structure, we store, for each set of alternatives, a triple containing the site at which the set occurs and the left and right boundary of the concatenated alternatives. In our example the triple corresponding to Figure 2 is (1, 2, 5). This information is then used to generate the *jump index array*. For each marker, it stores the markers at which a backward search continues (the targets). In a backward search, we match a pattern from right to left. If a site marker is reached, the matching continues at the right boundaries of the corresponding alternatives. If the left boundary of an alternative is reached, the matching continues at the corresponding site marker. In our example, we obtain the *jump index array* shown in Table 2. Because marker number 5 appears in *S* directly before the sentinel $, it will never be encountered in a backward search.

**Table 2:** The *jump index array* for the example from Figure 2.

| marker | 1 | 2 | 3 | 4 | 5 |
| --- | --- | --- | --- | --- | --- |
| site marker | true | false | false | false | false |
| jump target | 3,4,5 | 1 | 1 | 1 | - |

#### 2.2.1 Nested variations

The *jump index* can deal with nested variations because it is possible to insert a site marker into an alternative.

#### 2.2.2 Encoding copy number variations

Copy number variations (CNVs) seem to be the most frequent structural variants in the current data. A CNV specifies the possible number of copies of a substring at a certain position. To illustrate our encoding, suppose that *ω* is a substring that occurs once at position *p*, but in a genetic variant it occurs twice at *p*. In our encoding, we would insert two site markers at the variant site *p*. Both markers trigger a backward search for *ω*, which is continued at the position immediately before the marker that triggered the search. In this way, we take care of the two occurrences of *ω*. To take the single occurrence of *ω* into account, we add a ‘jump’ from the marker at position *p* to itself. This is equal to ignoring the marker.

In general, if *c* is the maximum copy count of *ω*, we add *c* site markers. For each other observed copy count *ĉ*, we add a ‘jump’ from the marker (*r* + *c* − *ĉ* − 1) to marker *r*, where *r* is the rank of the first site marker of this variation.

#### 2.2.3 Encoding other structural variants

Transpositions and non-tandem duplications of a string *ω* can be encoded similarly. The difference to CNVs is that *ω* may occur at several positions *p*_1_, *p*_2_,… in the sequence. Site markers for such a variant must be inserted at those positions. For each site marker a ‘jump’ to itself is added, so that a backward search for *ω* may ignore the marker. Inversions are encoded in the straightforward way, in which the set of alternatives consists of the reverse complemented string.

#### 2.2.4 Transforming the information

Till now, we identified a marker by its rank in the *jump index string*. However, a backward search is based on the BWT of the *jump index string*. The rank of a marker in the *jump index string* differs from its rank in the BWT. For example, the fifth marker in the *jump index string* is equal to the first marker in the BWT of Table 1. To make use of the data in the *jump index array*, we need to calculate the permutation that maps the rank of a marker in the *jump index string* to its rank in the BWT. We note that the order of the markers in the BWT coincides with their order in F, where F[*i*] = S[SA[*i*]] (see Table 1 for an example). Clearly, the ranks of the markers in the *jump index string* have the same order as their positions in that string. Moreover, if there are *m* occurrences of # in the *jump index string*, then the suffix array interval [2..*m*] contains all the positions at which markers occur in the *jump index string*. It follows as a consequence that the problem of calculating the permutation can be solved by sorting the numbers in SA[2..*m*]; see Table 3. To be precise, for each marker we store a pair (position in the *jump index string*, rank in the BWT) and sort the pairs by the positions. Afterwards, we replace positions by their ranks in the *jump index string* and obtain the desired permutation (its inverse permutation can easily be computed). We use the permutation to transform the jump index array. The information stored after this transformation is shown in Table 4.

**Table 3:** Calculation of the permutation from rank in *jump index string* to rank in BWT.

|  |  |  |  |  |  |
| --- | --- | --- | --- | --- | --- |
| position in *jis* | 18 | 4 | 12 | 16 | 8 |
| rank in BWT | 1 | 2 | 3 | 4 | 5 |
| ↓ sort by position in *jis* |  |  |  |  |  |
| position in *jis* | 4 | 8 | 12 | 16 | 18 |
| rank in BWT | 2 | 5 | 3 | 4 | 1 |
| ↓ replace positions by ranks |  |  |  |  |  |
| rank in *jis* | 1 | 2 | 3 | 4 | 5 |
| rank in BWT | 2 | 5 | 3 | 4 | 1 |

**Table 4:** The *jump index array* after transformation.

|  |  |  |  |  |  |
| --- | --- | --- | --- | --- | --- |
| marker | 1 | 2 | 3 | 4 | 5 |
| site marker | false | true | false | false | false |
| jump target | - | 1,3,4 | 2 | 2 | 2 |

## 3 Exact Matching Algorithm

Algorithm 1 shows our backward search algorithm in pseudo-code. In each iteration of the while-loop, a backward search step is made for all intervals. If such a step, applied to an interval *iv*, returns an interval [*l..r*] with *l* > *r*, then the search failed and the interval iv is deleted. In the algorithm, an interval is actually a triple, consisting of the left and right boundary of a suffix array interval and a return stack. The stack is needed to handle jumps. The two procedures *snp_handling()* and *jump_handling()* (Algorithms 2 and 3) add new intervals to the set of intervals.

Algorithm 2 performs a search step for all IUPAC letters that match the current character, but are not identical to it.

Algorithm 3 searches for markers in the current intervals. Each found marker is processed in the for-loop at line 4. If the marker is a site marker, then a singleton interval is added to the set of intervals for each right boundary of the alternatives in the corresponding set. The latter can be found in the *jump index array*. For each added interval, the interval of the site marker is pushed onto its return stack. This has the effect that, after processing the alternative, the algorithm ‘jumps’ back to this variant site. More precisely, if the current marker is not a site marker, then the algorithm has reached the end of an alternative. If the return stack of the alternative is not empty, then its top element is the corresponding variant site. If the return stack is empty, then the search started inside an alternative. In this case, the singleton intervals of all variant sites, at which this alternative can occur, are added to the set of intervals.

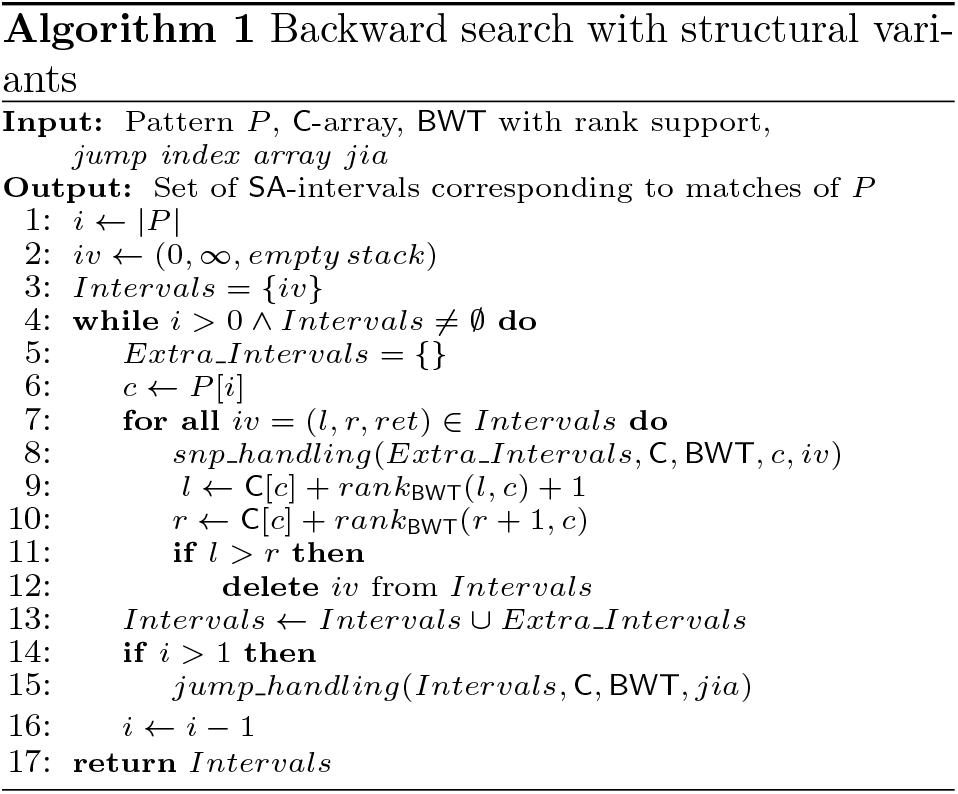

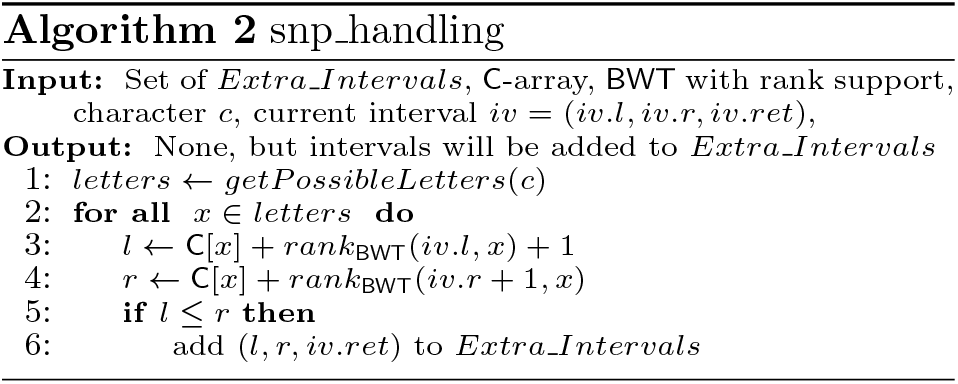

### 3.1 Accelerating the search with a *k-mer-index*

At the beginning of a search, the intervals are relatively large. Hence they may contain many markers. For each site marker, the procedure *jump_handling()* adds at least one interval to the set of intervals. A a result, the number of intervals increases dramatically in the first search steps. Because most of the intervals are deleted after a few more steps, the number of intervals then decreases rapidly. In other words, most of the computation time is spent in the first few iterations. To speed up the search, *jisearch* provides an option to use a *k-mer-index* (*k* is a parameter that can be set by the user, *k* = 10 is recommended). More precisely, it precalculates the suffix array interval of each of the 4^*k*^ *k*-mers (strings of length *k* on the DNA-alphabet); if the *k*-mer is not a substring of *S*, it stores the empty interval. The calculation of the *k-mer-index* is done iteratively: the suffix array intervals of all *q*-mers are computed based on suffix array intervals of all *q* − 1-mers, where 1 ≤ *q* ≤ *k*. In a backward search for pattern *P, jisearch* looks up the suffix array interval of the *k*-mer suffix of *P* in the *k-mer-index* and then the search continues with *P* [1..|*P*| − *k*]. A similar technique was also used by Huang *et al*. (2013) and Maciuca *et al*. (2016).

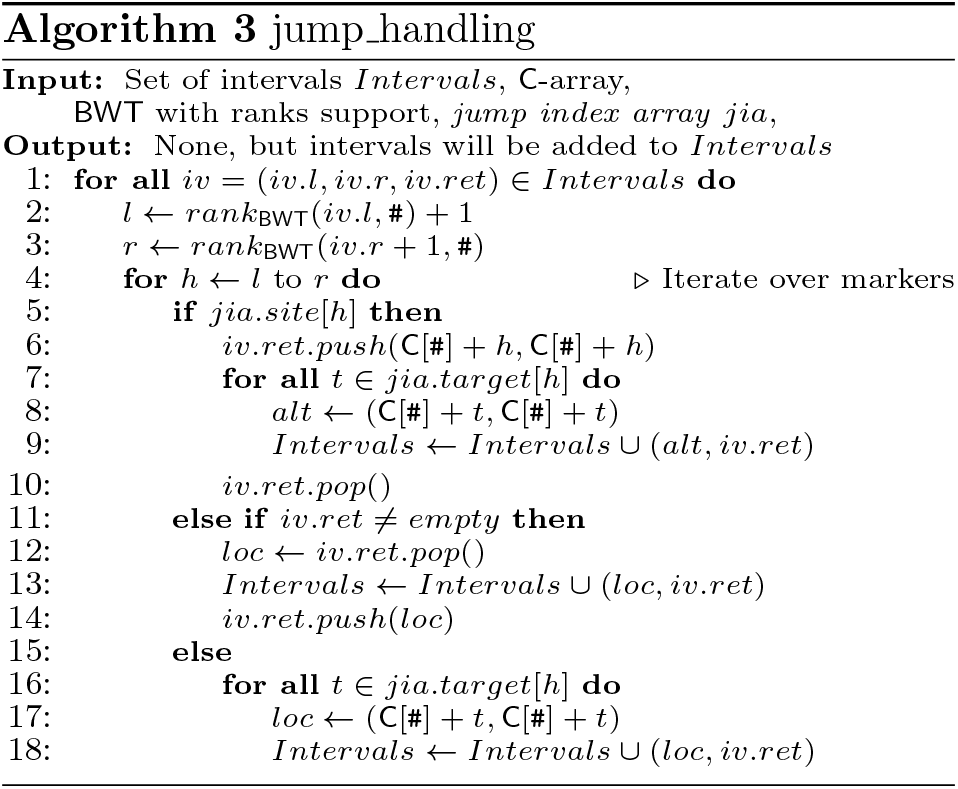

### 3.2 Accelerating the search for singleton intervals

The procedure *jump_handling()* generates many single-ton intervals. A singleton interval represents just one position in the *jump index string*. In this case, it is more efficient to search the pattern in the *jump index string* (character-by-character) than using a BWT-based backward search (because string access is much faster than rank queries). To implement the string search, we need a second set called position set. In this set, we store the position SA[C[#] + *t*] instead of the interval [C[#] + *t*..C[#] + *t*]. The character-by-character search starts at position SA[C[#] + *t*] in the *jump index string* and if it is done from right to left (i.e. backwards), then it can be integrated into the overall search algorithm. If the current character in the *jump index string* is a marker, we need to know its rank in the BWT to load the jump targets. Therefore, we store a map that maps the position of a marker in the *jump index string* to its rank in the BWT. In fact, we used this map already; see Section 2.2.4.

### 3.3 Output

The matching program of *jisearch* prints information about the number of found reads and stores a binary representation of the interval and position sets. To obtain detailed information, a second program can be used that processes the binaries. It prints the name of each pattern *P* and for each occurrence of *P* in the sequence it prints the following information: the chromosome name, the position at which *P* occurs in the reference sequence, the corresponding position in the *jump index string*, and if the match includes variations, a list containing the variation IDs and the used alternatives.

If the search for pattern *P* results in a suffix array interval [*l..r*], the positions of the occurrences of *P* in the *jump index string* are SA[*i*], where *l* ≤ *i* ≤ *r*. For each position, a string forward search is executed that logs the ID and the chosen alternative for each jump. The corresponding position in the reference sequence and the chromosome name are queried in a separate data structure. In principle, the same information could be calculated during the backward search (i.e. within the program *jisearch*). However, since we do not know in advance for which intervals the search succeeds, this must be done for all the suffix array intervals that are created during the search. By contrast, the additional forward search is solely applied to the (relatively few) positions at which the pattern matches. Our experiments confirmed that it is advantageous to use the additional forward search.

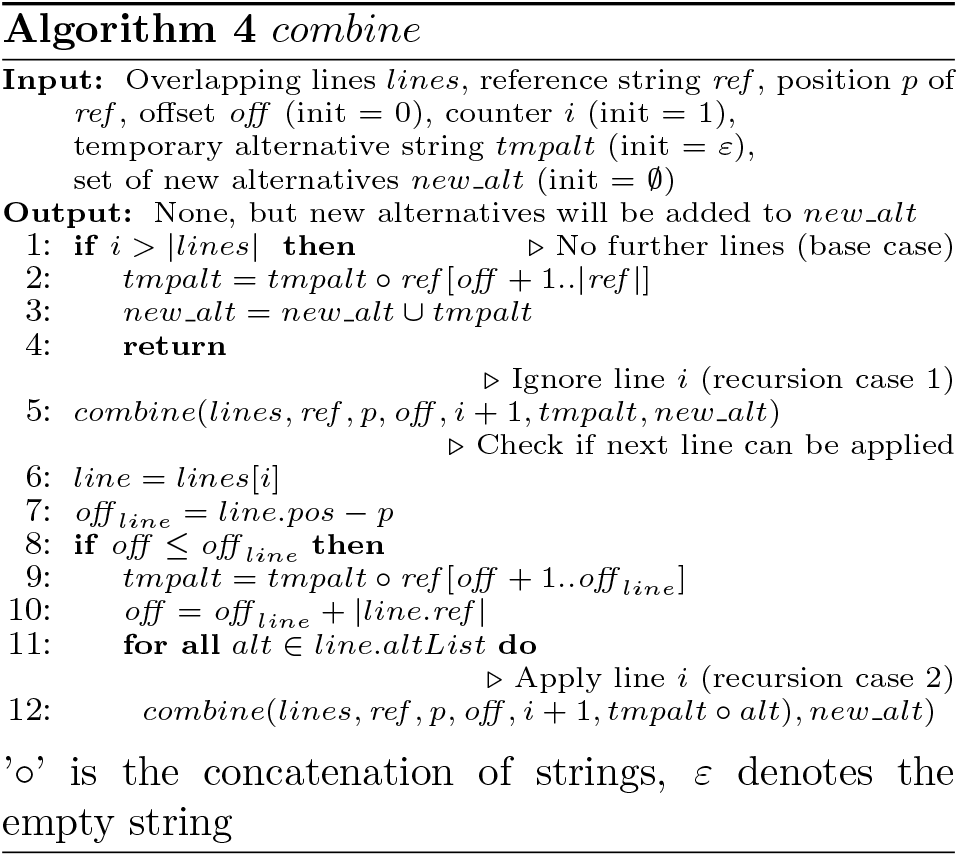

## 4 Implementation and experiments

Our software tool constructs the *jump index string* from a reference genome in the form of a FASTA-file and a list of variations in the form of a VCF-file. For the DNA sequence of a chromosome, the *jump index string* is built as follows. Let *p* be a position in the chromosome that occurs in the list of variations and let *ω* be the substring that starts at *p* and for which alternatives are listed. Then *ω* is cut out of the DNA sequence and replaced with a marker #. The listed alternatives and *ω* are appended at the end of the string, each separated by #; see Figure 2.

The test data was taken from the 1000 Genomes Phase II (hs37d5) sequence^1^ and the corresponding variation list.^2^ We constructed the search index for the first human chromosome, which has approximately 250 million base pairs. The variation file for this chromosome contains about 6.5 million entries, 96% of which represent SNPs. Consecutive lines in the VCF-file can have overlapping references, resulting in overlapping variants, and we have to deal with this problem. In the following, it is shown how such lines can be combined to a new line that describes all variants. As an example, consider the reference string ‘TTAA’ and the three variations: (1:TTAA:T), (2:T:A), (3:A:AT). Each triple consists of a position, a reference, and a list of alternatives. The first triple describes the deletion of ‘TAA’ at position 1, the second a SNP at position 2, and the third an insertion of ‘T’ after position 3. We will combine these to (1:TWAA:T,TWATA). In this triple, the SNP is represented by its IUPAC code letter ‘W’ and there are two alternatives: the first one is the deletion and the second one is the insertion. To calculate the new triple, we first determine the shortest substring that covers all references of the triples (lines) under consideration. Then the letters at the position of a SNP are replaced by the corresponding IUPAC code letter. In our example, the new reference *ref* is ‘TWAA’. After that, Algorithm 4 is called with the lines that do not correspond to SNPs, the new reference *ref* and the starting position *p*. The algorithm runs recursively until all lines have been processed. Each line has the components *pos, ref*, and *altList*. In each recursive step, it is checked whether the line can be applied or not, i.e., whether its alternatives can be added or not. The algorithm has two cases, the first ignores the line and the second applies the line if possible. So each possible combination of ignoring or applying the lines is generated and stored in the set *new_alt*. The new alternative of ignoring all lines is identical to the reference. The string *tmpalt* is used to gradually build a new alternative by appending parts of *ref* or the alternatives of a line. A line cannot be applied if parts of its reference were changed by a previous line. To decide whether this is the case, the parameter *off* stores the (relative) position of the last letter of *ref* that was changed. If *off* is larger than the offset of the line (*off_line_*), the reference of the line was changed previously and the line cannot be applied.

Our index data structure is constructed by using the library SDSL (Gog *et al*., 2014). It uses a wavelet tree that supports rank-queries on the BWT and a sampled suffix array to reduce the memory consumption; see e.g. (Ohlebusch, 2013) for details. Moreover, the *jump index array* is compressed by a technique that uses bit vectors in conjunction with constant-time rank-queries; see Table 5. This works as follows. For each marker, the *jump index array* stores the information whether the marker is a site marker or separator and at which markers the backward search continues (targets). The marker type is stored in the bit vector *site*. That is, site[*i*] = 1 if and only if the marker of rank *i* is a site marker. Naively, the targets can be stored in a vector *targets* of vectors, so that *targets*[*i*] is the vector containing the targets of marker *i*. The target vectors can have arbitrary size, but the size is mostly one or two. We compress this representation by storing the target vectors with size 1 or 2 in integer vectors *v*1 and *v*2, respectively. The target vectors with other sizes are stored in the vector of vectors *v*3. Let *m* be the number of markers and let *o* and *t* be the number of markers with one and two targets, respectively. We define *bv*1 as a bit vector of length *m* with bv1[*i*] = 1 if and only if |targets[*i*]| = 1. We define *bv*2 as a bit vector of length *m − o* with *bv*2[*j*] = 1 if and only if |*targets*[*i*]| = 2, where *i* is the position of the *j*-th 0 in *bv*1. Note that there are *o* ones in *bv*1 and *t* ones in *bv*2. Furthermore, |*v*1| = *o*, |*v*2| = 2*t*, and |*v*3| = *m − o − t*. The five new vectors need much less space than the original vector of vectors. Table 5 shows the compression of the target vector of our example. For fast access to the original targets, the bit vectors *bv*1 and *bv*2 are preprocessed so that constant-time rank-queries are possible. If the targets of marker *i* should be loaded, *r*_1_ = *rank*_1_(*bv*1, *i*) is calculated. If *bv*1[*i*] = 1, then *i* has just the one target *v*1[*r*_1_]. Otherwise, *r*_2_ = *rank*_1_(*bv*2, *i − r*_1_) is calculated. If *bv*2[*i − r*_1_] = 1, then *i* has the two targets *v*2[2*r*_2_] and *v*2[2*r*_2_ + 1]. Otherwise, the target vector can be found in *v*3[*i − r*_1_ − *r*_2_].

**Table 5:** Compressed target vector of the *jump index array* in Table 2

|  |  |  |  |  |  |
| --- | --- | --- | --- | --- | --- |
| *bv1* | 0 | 0 | 1 | 1 | 1 |
| *bv2* | 0 | 0 |  |  |  |
| *v1* | 2 | 2 | 2 |  |  |
| *v2* |  |  |  |  |  |
| *v3* | $\langle \rangle$ | $\langle 1,3,4 \rangle$ | | | |

We evaluated the performance of *jisearch* and compared it to *gramtools*^3^ and *BWBBLE*. Each of the tools takes the above-mentioned test data (the human chromosome 1 and the corresponding VCF-file) as input and generates its internal data structures. If a tool cannot deal with a certain type of genetic variant in a VCF-line, it simply skips the line.

All experiments were conducted on a Ubuntu 16.04.4 LTS system with two 16-core Intel^®^ Xeon^®^ E5-2698 v3 processors and 256 GB RAM. Figure 3 shows the size of the interval set during the first search steps. As one can see, the number of intervals increases dramatically at the very beginning and then decreases exponentially (note the semi-log scale). This demonstrates why the first search steps need most of the computation time and why a *k-mer-index* is so useful. For a comparison with the other programs, we searched for exact occurrrences of the first 100k reads of the file SRR062634^4^ that do not contain the letter ‘N’. The programs *jisearch* and *gramtools* also search for the reverse complement of a read, so one read causes two searches. In *BWBBLE* the sequence is concatenated with its reverse complement, so there is no need to execute an additional search.

**Figure 3:**
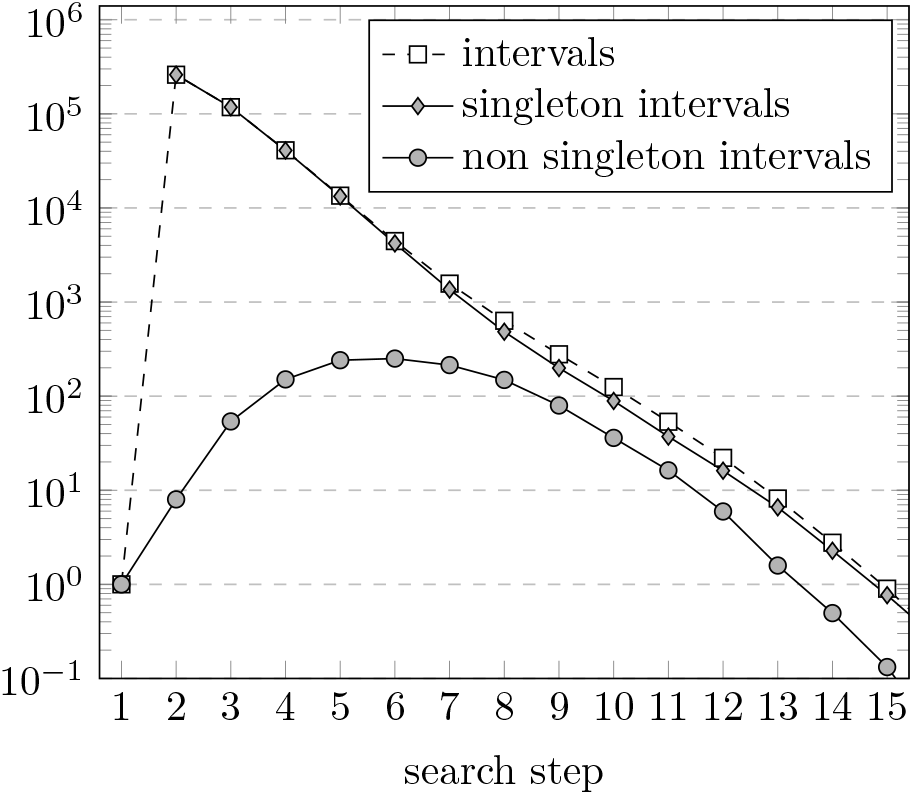
10k randomly generated patterns were searched for in the first human chromosome. The figure shows the average size of the interval set at the beginnings of the first 15 search steps and the average number of singleton and non-singleton intervals.

We tested *jisearch* with a *k-mer-index* for *k* = 5, and *k* = 10 (gramtools uses *k* = 5). Figure 4 compares the tools with respect to index construction time, index size, and the mapping speed. On one hand, the construction time and index size of a *k-mer-index* increases with *k*. On the other hand, a larger *k-mer-index* accelerates the mapping speed significantly. As one can see, the index of *jisearch* needs less space and supports faster search than the index of *gramtools*. The reason is probably the enormous alphabet size in *gramtools*. The string generated by *BWBBLE* is much longer than the strings generated by the other programs because it includes the reverse complement of the sequence and a ‘padding’ for each variant. This and probably the fact that Huang *et al*. (2013) did not use the SDSL leads to a bigger *FM-index. BWBBLE* reaches a high mapping speed without using a *k-mer-index*.^5^ This can be attributed to the fact that their backward search does not need to follow different edges in the pan genome reference graph. As indicated by the title of the article (Huang *et al*., 2013), the method only works well for short reads. By contrast, *jisearch* and *gramtools* are able to search for reads of arbitrary length. The percentage of mapped reads is lower that 7% for all programs. This is probably because the chromosome 1 represents about 8% of the human genome. Our program maps (a few) more reads than the others. *BWBBLE* has problems in mapping reads with variations that are too near to each other and *gramtools* ignores a variation completely if its starting position is covered by another variation.

**Figure 4:**
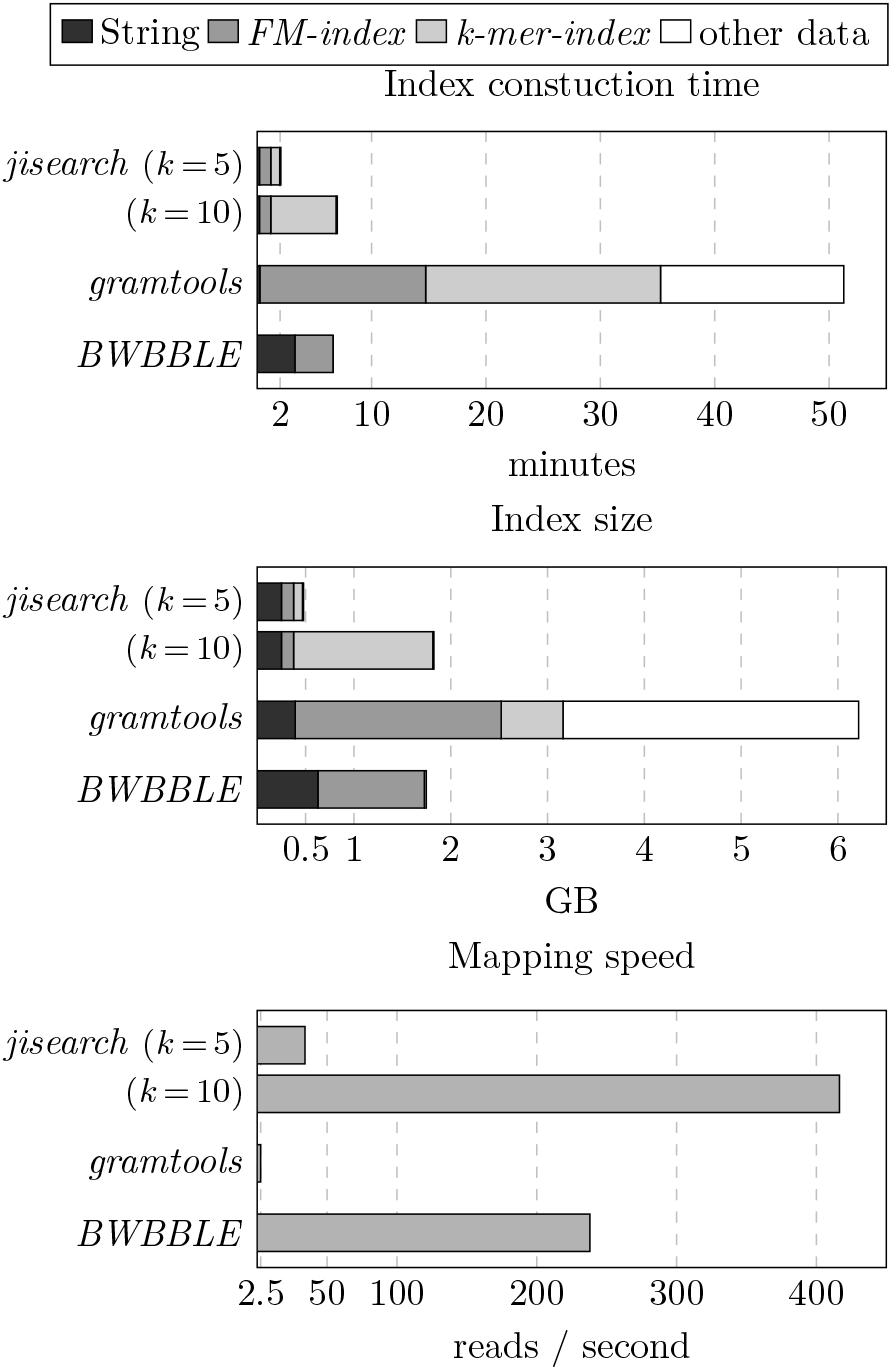
Comparison of the indexes generated by the tools. The first diagram displays the time for the index construction. The second diagram shows the size of the index on disc. All tools generate a string and calculate its *FM-index*, but only *jisearch* and *gramtools* calculate a *k-mer-index*. ‘Other data’ refers to additional data that a tool might need in a specific application. The third diagram shows the mapping speed in reads per second.

## 5 Discussion

We have presented a new method to encode genetic variation in a Burrows-Wheeler transform, which extends the work of Huang *et al*. (2013) and Maciuca *et al*. (2016) in several aspects. Apart from SNPs and indels, this method allows to encode larger structural variants such as large deletions, inversions, copy-number variation, duplications, and transpositions. (Since the current data lacks non-tandem duplications and transpositions, these are not yet supported by our implementation.) Moreover, the method can deal with nested variations. In contrast to gramtools, our method uses only one extra symbol and therefore avoids the disadvantages of an increased alphabet size. As gramtools (Maciuca *et al*., 2016), the software *jisearch* could be used to “support the inference of the closest mosaic of previously known sequences to the genome(s) under analysis.” Our future goal, however, is to extend the exact matching algorithm in such a way that it supports inexact matching. It is unclear whether the techniques used in the software tool *BWBBLE* (Huang *et al*., 2013) can be adapted to our more general setting; it might be the case that different methods have to be developed to efficiently cope with larger structural variants in read mapping.

## Footnotes

1 FASTA: ftp://ftp.1000genomes.ebi.ac.uk/vol1/ftp/technical/reference/phase2_reference_assembly_sequence/

2 VCF: ftp://ftp.1000genomes.ebi.ac.uk/vol1/ftp/release/20130502/

3 Version 0.5.0 from 09/2018, commit hash ad47bb6…

4 FASTQ: ftp://ftp.1000genomes.ebi.ac.uk/vol1/ftp/phase3/data/HG00096/sequence_read/

5 Although the possible usage of a *k-mer-index* is described in (Huang *et al*., 2013), it seems that the software *BWBBLE* does not use it at all.

